# Spatial Ecology of Jaguars in a Coastal Amazon Island Ecosystem

**DOI:** 10.1101/2025.07.10.664043

**Authors:** Leandro Silveira, Anah T. A. Jácomo, Tiago J. Silveira, Gediendson R. de Araujo, Renato A. Moreira, Francesca B. L. Palmeira, Marcia de A. M. M. Ferraz, Michael J. Noonan

## Abstract

Understanding the spatial ecology of large carnivores is essential for effective conservation, particularly in insular or fragmented systems where movement is constrained. Jaguars are particularly important to study, as they are opportunistic predators whose diet and behaviour shifts flexibly with local prey availability. We investigated the movement patterns, home range dynamics, and social interactions of jaguars (*Panthera onca*) on Maracá-Jipioca Island, a pair of small, seasonally flooded islands, in a low-disturbance reserve in the Brazilian Amazon. Using GPS collars, we tracked seven individuals and found that males had significantly larger home ranges (mean: 77.8 km²) than females (mean: 19.5 km²). Despite the island’s limited area, home-range overlap was low, and direct encounters were rare. This mirrored the territorial behaviour seen in more expansive mainland populations, but contradicted patterns seen in other populations that specialised on aquatic prey. Jaguars on Maracá-Jipioca moved an average of 7.3 km/day, with no significant differences across sex, age, nor body size. However, movement speed and probability of activity varied significantly across habitats: individuals moved faster in grasslands, wetlands, and water, and more slowly in forests and mangroves. Animals also spent more time resting while in forests and mangroves as compared to grasslands, water, and wetlands. In contrast to the predominantly nocturnal habits reported in continental populations, jaguar activity on Maracá-Jipioca peaked during daylight hours, especially among females. Jaguars showed a marked preference for coastal areas but did not appear to adjust habitat use in response to tidal variation in sea level. Collectively, these findings suggest that the spatial ecology of Maracá-Jipioca jaguars is comprised of typical and novel behaviours, shaped by both the species evolutionary history, and the island’s unique ecological context. These findings highlight the species’ behavioural flexibility and offer rare insights into how a large carnivore persists in an insular Amazonian landscape.

## Introduction

Jaguars (*Panthera onca*) are the largest felid in the Americas and the third-largest in the world [1]. Historically ranging from as far north as the southwestern United States to central Argentina [1–3] in the south, the species demonstrates a remarkable amount of ecological plasticity, occupying environments as diverse as dense forests and extensive wetlands to xeric desert ecosystems [4]. In addition to being habitat generalists, jaguars are opportunistic predators whose diet shifts flexibly with local prey availability [5–8]. They have been documented feeding on >85 species [9], ranging from large mammals and cattle [10] to small rodents [11], reptiles [7], and fish [12]. As a top predator, jaguars can shape the ecosystems they occupy through top-down regulation [1,13] and competitive exclusion [14]. However, jaguars’ own ecology is also heavily impacted by their feeding ecology. For instance, their body mass ranges from ∼56kg (males), ∼41kg (females) in Central America to ∼105kg (males), ∼77 kg (females) in the Pantanal, and is thought to be governed by their hunting decisions [8]. Similarly, while they tend towards solitary behaviour with social interactions limited to mating and territorial disputes throughout much of their range [15], populations that rely on aquatic prey bases have been found to exhibit cooperative fishing, co-traveling, and play [12]. Given jaguars’ high degree of morphological, dietary, and behaviour plasticity, alongside their keystone role in shaping the ecosystems they occupy [13], it is important to understand how jaguar ecology changes throughout their range.

Although jaguars are currently listed as near threatened by the IUCN [16], and occupy only about half of their historical distribution, due primarily to habitat loss, human conflict, and prey decline [17–20], they still remain widely distributed across various biomes in Brazil, including the Amazon, which harbours the highest remaining densities and thus represents a stronghold for their conservation. While an extensive body of research has elucidated the species’ ecology across a range of continental environments [21], their habits within insular, mangrove-dominated ecosystems have remained largely uninvestigated. Within the Amazon biome, Maracá-Jipioca Ecological Station (ESEC-MJ), a pair of insular landmasses off the Amapá coast, presents a unique environment where jaguars thrive despite spatial constraints. Despite their small size (ca. 600 km^2^), these islands offer a mosaic of habitats including mangroves, flooded grasslands, and tidal wetlands that support a high abundance of aquatic and semi-aquatic prey [22]. The isolation of the islands, combined with their protected status and prey availability, may create a rare natural refuge where jaguar populations persist with minimal anthropogenic interference [23]. Previous studies in the region have documented exceptionally high jaguar densities (up to 6.7 individuals/100 km²) and unusual behaviours, including diurnal activity and foraging on marine fauna [23,24].

In this study, we investigated the movement patterns of jaguars inhabiting the Maracá-Jipioca islands using GPS collars deployed on seven adult individuals. Understanding movement patterns is essential for conservation planning, as it informs habitat use, territoriality, and resource exploitation strategies [21,25]. In spatially restricted systems like Maracá-Jipioca, movement data also reveal how large predators cope with physical and ecological constraints, including the trade-offs between prey search, territorial boundaries, and reproductive opportunities [26–28]. Such information is critical for evaluating the long-term viability of island populations and the ecological processes they maintain. Building on previous research documenting this population’s high density and unique dietary habits [22,24], we sought to understand how these large predators navigate a spatially constrained and hydrologically dynamic environment. Specifically, we estimated these individuals’ home-range areas, movement rates, and their circadian rhythms. Given that aquatic prey bases have been previously shown to impact their social behaviour [29], we also investigated patterns of home-range overlap and their encounter rates, with attention to potential differences between sexes. Lastly, we analysed their habitat preferences, and the extent to which these were impacted by the tidal nature of this island ecosystem. By characterising their movement ecology, this study provides key insights into how jaguars persist in isolated islands and offers critical information to guide conservation and management strategies within this protected Amazonian refuge.

## Methods

### Sampling area

Jaguar movement data were collected from animals on Maracá-Jipioca Ecological Station (ESEC-MJ), a federal protected area located off the coast of Amapá State in the northeastern Brazilian Amazon (02°01′13″N, 50°30′20″W). The protected area spans approximately 60,000 hectares, comprising two main continental-origin islands: Maracá and Jipioca, separated from the mainland by estuarine channels approximately 6–10 km wide. The islands were formed by erosional and sedimentary processes over the past 10,000 years [30,31], and are strongly influenced by the dynamics of the Amazon River delta and the Atlantic tide, which produces daily flooding across low-lying areas.

ESEC-MJ lies within the humid tropical zone and experiences a tropical monsoon climate (Am, Köppen classification), with annual rainfall ranging from 2,300 to 2,800 mm and average temperatures between 25–27°C. The region has minimal elevational variation, with a mean altitude of approximately 3 meters above sea level, making it highly susceptible to tidal and seasonal flooding. The landscape consists of a mosaic of interconnected wetlands, including mangroves (Rhizophora and Avicennia), flooded grasslands (dominated by Juncus spp.), freshwater lagoons, estuarine channels, bamboo thickets (Guadua spp.), and patches of coastal scrub and low forest. According to the 2023 MapBiomas land classification maps (http://mapbiomas.org), Maracá and Jipioca islands are composed primarily of ca. 26.5% forest, 18.8% mangrove, 50% water and wetland, and 4.6% grassland.

This unique habitat configuration supports a high biomass of aquatic and semi-aquatic prey, including fish, turtles, birds, and caimans, which jaguars regularly exploit. Human presence on the islands is minimal due to the station’s protected status, with no permanent settlements, roads, or agriculture, making it one of the few undisturbed coastal island systems harbouring a viable large carnivore population in the Amazon.

### Animals capture and GPS data acquisition

Between 2019-10-04 until 2025-01-24, seven jaguars were captured using either box traps, foot snares, or trained hounds. Captures via box traps employed wooden box traps (90 × 90 × 200 cm) with metal doors that were baited with fresh fish. Traps were strategically placed in shaded areas along the western edge of the island in locations with documented prior jaguar activity [22,24]. Traps were inspected twice daily, during early morning and late afternoon. In addition to box traps, foot snares were deployed on trails previously assessed for soil firmness to ensure secure anchoring. Snare installation followed procedures outlined by de Araujo et al. 2021 [32], incorporating a torsion-spring-powered thrower, a robust anchoring system using rebar stakes, and a VHF telemetry-based remote monitoring system allowing checks at maximum one-hour intervals. Transmitter activation occurred via magnetic switch detachment, facilitating rapid response upon capture. Trained hounds were also used to track fresh jaguar scent trails, consistent with McBride & McBride (2007) [33]. The hounds were used to locate and bay the animals, enabling remote chemical immobilization via a dart projector rifle. The team was prepared to manage arboreal containment situations, including deploying safety nets to mitigate risks during tree descents.

In the first campaign (2019), three individuals were captured; in the second (2023), two individuals, including the recapture of a female (named Iemanja); and in the third (2024), three additional individuals. Captured individuals (n=7) were chemically immobilized by intramuscular administration of medetomidine (0.1 mg/kg) and ketamine (5 mg/kg). Additionally, biological samples were collected, biometric measurements were recorded, and the animals were fitted with GPS collars (Telonics® TGW-4577-4 and Iridium Terrestrial Systems). The GPS collars were set to record locations at either 30, or 60 minute intervals (Table S1). Once all procedures were completed, anesthesia was reversed using Atipamezole (0.25 mg/kg, Precision Pharmacy) or Yohimbine (0.3 mg/kg). Veterinary supervision was maintained throughout the entire process to ensure animal welfare. The capture and handling protocols complied with ethical standards and received approval from the Chico Mendes Institute for Biodiversity Conservation, under license number 63094, and the Ethics Committee for Animal Use at the University of Mato Grosso do Sul, under license number n° 1.221/2022. All procedures strictly adhered to the ethical guidelines of the American Society of Mammalogists [34].

### Animal movement analysis

#### Movement data pre-processing and modelling

Before analysis, we performed a data cleaning process to remove any outliers using the methods implemented in the `R` package `ctmm` [35]. For each individual dataset, we first removed entries without coordinates, or with duplicate timestamps for all animals. We then removed outliers based on distance from the median location, and the minimum speed required to explain each location’s displacement. Through this second process, we identified a total of 18 outliers, which represented <0.02% of the 68,934 GPS locations. For each of the collared jaguars, we then confirmed range-residency via variogram analysis (Fig. S1; [36]), fit a series of continuous-time movement models to the location data, using perturbative-hybrid residual maximum likelihood (pHREML; [37]), and identified the best model via small sample-sized corrected Akaike’s information criterion (AICc). The models were also validated via variogram analysis, and all were found to match the key features in the data (Fig. S1). The selected movement models formed the basis of all of our downstream movement analyses.

#### Home-range behaviour

After fitting the movement models, we estimated each animal’s home-range area as the polygon delimited by the 95% isopleth of the utilization distribution using autocorrelated kernel density estimation (AKDE; [38]). These home-range estimates were conditioned on the movement models’ autocorrelation structures, and we implemented the small sample size bias correction of Fleming & Calabrese (2017) [39], and the optimal weighting of Fleming et al. (2018) [40]. Because these animals lived on an island, we mitigated any spillover bias [41] by applying local kernel correction based on a spatial polygon of Maraca’s boundary during the kernel density estimation process [42]. To test for a difference in home-range areas between sexes, we used the meta-regression methods implemented in `ctmm` [43]. Because `ctmm` only allows for comparing categorical variables, we tested for correlations between home-range size and both age and weight via the meta-regression methods implemented in the `R` package `metafor` [44]. This allowed us to propagate uncertainty in the individual home-ranges estimated into our downstream analyses.

#### Home-range overlap and social behaviour

Because we were also interested in describing these jaguars’ social behaviour, we estimated the amount of home-range overlap between all pairs of individuals (i.e., dyads) that were monitored at the same time via the Bhattacharyya coefficient [45]. To determine whether sex was a factor underpinning the degree of pairwise overlap, we fit a generalized linear mixed model (GLMM) with a beta distribution and a logit link function to the home-range overlap estimates, with pairwise sex as a predictor variable via the `R` package `mgcv` [46]. This model was then compared to a similar model that excluded the pairwise sex predictor variable using a generalised likelihood ratio test.

While home-range overlap describes patterns in spatial structuring, it does not directly indicate whether individuals are likely to be in the same place at the same time (Winner et al., 2018). We therefore also estimated a proximity ratio for all dyads via the `ctmm` function `proximity`, whereby a proximity ratio of 1 is consistent with independent movement; values >1 indicate individuals are further apart on average than expected for independent movement and vice versa for values <1 (see also [47]). For each dyad we also estimated the Euclidean distance using the `ctmm` function `distances`, from which we estimated the number of encounter events using a 100 m distance threshold.

#### Movement speed and activity patterns

In order to investigate the activity and movement patterns of jaguars on Maracá, we estimated each animal’s mean daily movement speed (in km/day) as well as the instantaneous movement speed (in m/s) at each of the sampled locations using continuous-time speed and distance (CTSD) estimation [48]. To test for a difference in average movement speed between sexes, we again used the meta-regression methods implemented in `ctmm` [43]. We then used a pair of GLMs with Tweedie distributions and log link functions to test for correlations between movement speed and both body weight, and age.

To study patterns of activity, we applied k-means clustering with three centers on the instantaneous speed estimates to identify a minimum speed threshold at which a jaguar could be classified as resting. Based on this analysis, any location with an estimated movement speed <1.5x10^-5^m/s was subsequently classified as “resting”, whereas locations with speeds >1.5x10^-5^m/s were classified as “active”. We also identified the land class at each location using the 2023 MapBiomas land classification rasters (http://mapbiomas.org) and the methods implemented in the `R` package `terra` [49]. To study how jaguars’ activity patterns differed between habitats, we used a hierarchical GLM with a binomial distribution and logit link to predict how active vs. resting (i.e., 0,1) behaviour differed as a function of habitat. For this model, individual ID was included as a random intercept. A subsequent GLM was then fitted to all of the instantaneous speed estimates where an animal was classified as active, with habitat as the predictor variable. Here we used a Tweedie distribution and log link, and again included individual ID as a random intercept. Finally, to study jaguars’ circadian rhythms, we used a hierarchical GAM with a binomial distribution, logit link, and individual ID as a random intercept to predict active vs. resting behaviour as a function of time of day. Here time of day was modelled using a cyclical cubic regression spline, and we also tested for an interaction between sex and time of day via a constrained factor smooth.

#### Habitat use and selection

To study patterns of habitat selection, we fit a hierarchical Poisson point process [50] using the bam function from the `mgcv` package. We opted for a GAM owing to its ability to capture the complex, non-linear patterns of habitat use that animals often exhibit [51]. Fitting a Poisson point process in this way required information on the sampling window and quadrature points. Because these animals lived on a pair of adjacent islands, we used the coastlines to define the sampling window. The quadrature points serve to approximate the likelihood function of a Poisson point process through Monte-Carlo Markov chain-based integration [50,52]. We therefore sampled locations uniformly throughout the islands, which resulted in a location:quadrature ratio of ca. 78:1. Following Alston et al. (2023) [53], we used autocorrelation-informed weights obtained from the individual home-range estimates to correct for autocorrelation-induced bias. The final model was of the form:

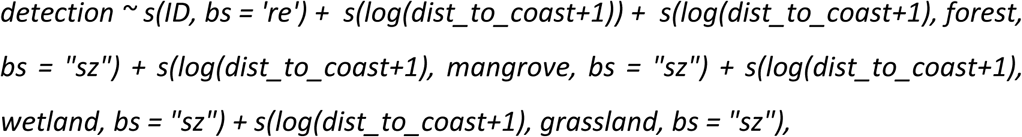

where detection represents whether the point was used by a jaguar or not (0, 1), and dist_to_coast was the distance to the coast (in m).

Finally, as these animals were observed to spend a substantial portion of their time close to the coast, we explored whether their habitat use was influenced by the tide. For this analysis, hourly data on the ocean height (masl) directly offshore of Maracá-Jipioca Island were obtained from the Copernicus Global Ocean Physics Analysis and Forecast [54]. We then used a GLM with a Tweedie distribution and log link function to assess whether individuals adjusted their distance to the coast based on the sea level.

All analyses were performed in `R` (ver. 4.3.3; R Core Team) [55], and the code required to reproduce all of the analyses is openly available in the GitHub repository at: https://github.com/QuantitativeEcologyLab/maraca_jaguars.

## Results

### Home-range behaviour

The average home-range size of jaguars on Maracá-Jipioca Island was 51.9 km^2^ (95% CI: 27.7 – 88.6 km^2^), ranging from 16.8 km^2^ (95% CI: 14.2 – 19.4 km^2^) to 128.8 km^2^ (95% CI: 73.9 – 198.8 km^2^). Males had a mean home-range size of 77.8 km^2^ (95% CI: 64.5 – 92.3 km^2^), versus 19.5 km^2^ (95% CI: 17.5 – 21.5 km^2^) for females. We estimated the male/female ratio of home-range areas to be 4.01 (3.23–4.88), which excludes 1, indicating males had significantly larger home-ranges than females (Fig. 1C). While there were differences in home-range size between the sexes, there were no significant relationships with neither weight (β = 0.001; 95% CI: -0.001 – 0.003; Fig. 1D), nor age (β = -0.008; 95% CI: -0.028 – 0.011; Fig. 1E).

**Figure 1.**
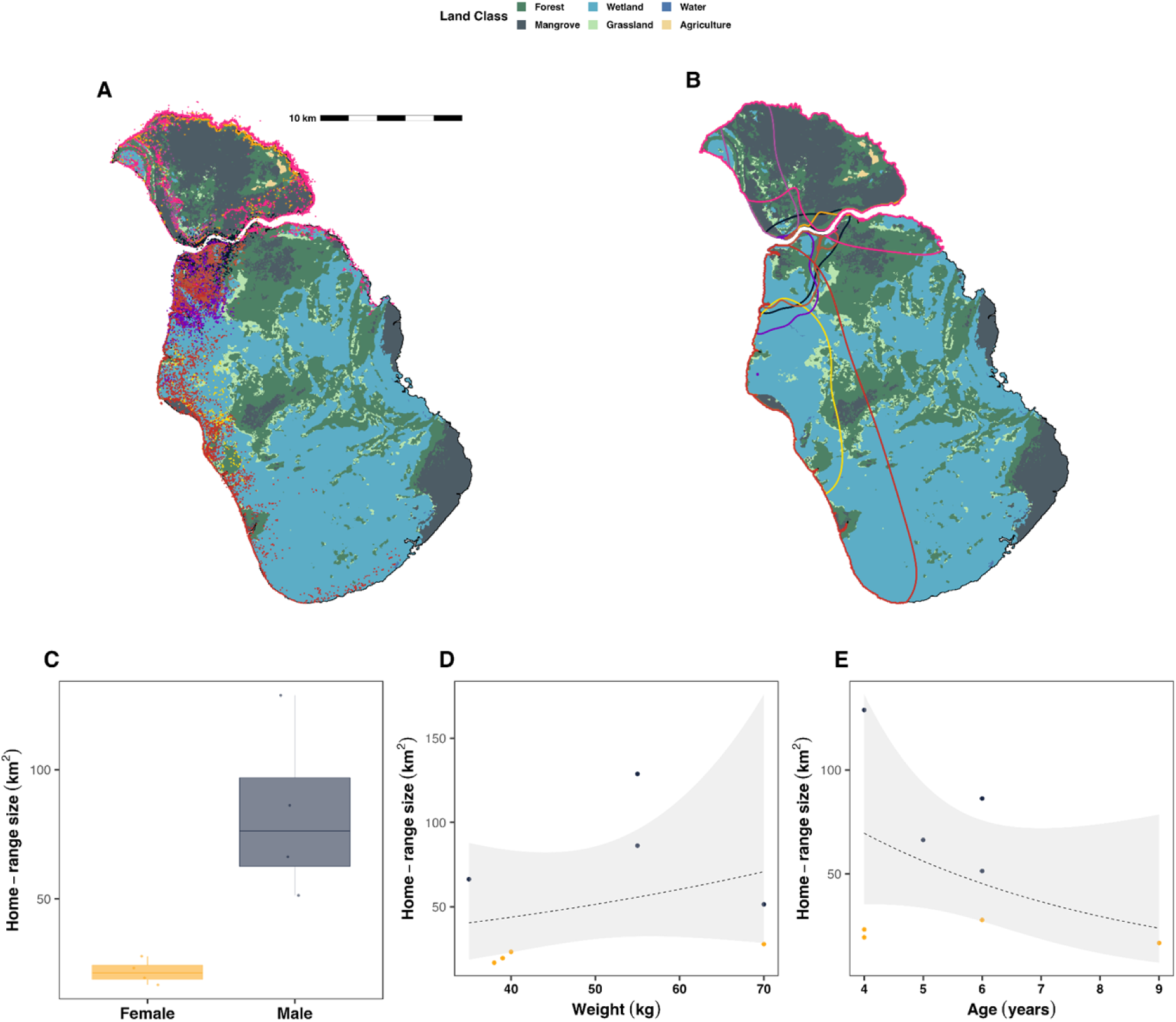
Jaguar space use and home-range behaviour. Maps depicting the (A) GPS locations, and (B) home ranges of the 7 monitored animals. In both panels the colours of the points and contours correspond to individual animals. The boxplot in (C) shows how home-range size differs between the sexes, while the scatterplots show home-range size versus (D) weight, and (E) age.

### Home-range overlap and social behaviour

The average home-range overlap for the five animals monitored during the same time interval was 0.24, but ranged from 0.016 between a pair of males (ID696469B and ID696490B) to 0.88 between another pair of males (ID696490B and Iranildo). There was no evidence of significant differences in pairwise overlap between any of the sexes (Λ_GLRT_ = 0.014, P = 1.0; Fig 2A). Furthermore, all but one of the proximity ratios overlapped 1, indicating no significant avoidance, nor attraction behaviour (Fig. 2B). Encounters between individuals were rare, with only an estimated 33 encounter events over the entire tracking period (Fig. 2C). Of these, 30 were encounters between two male-female pairs (Iranildo and Iemanja, and ID696469B and ID717047B), and 3 were encounters between Iranildo and ID696490B, the pair of males with the greatest amount of home-range overlap.

**Figure 2.**
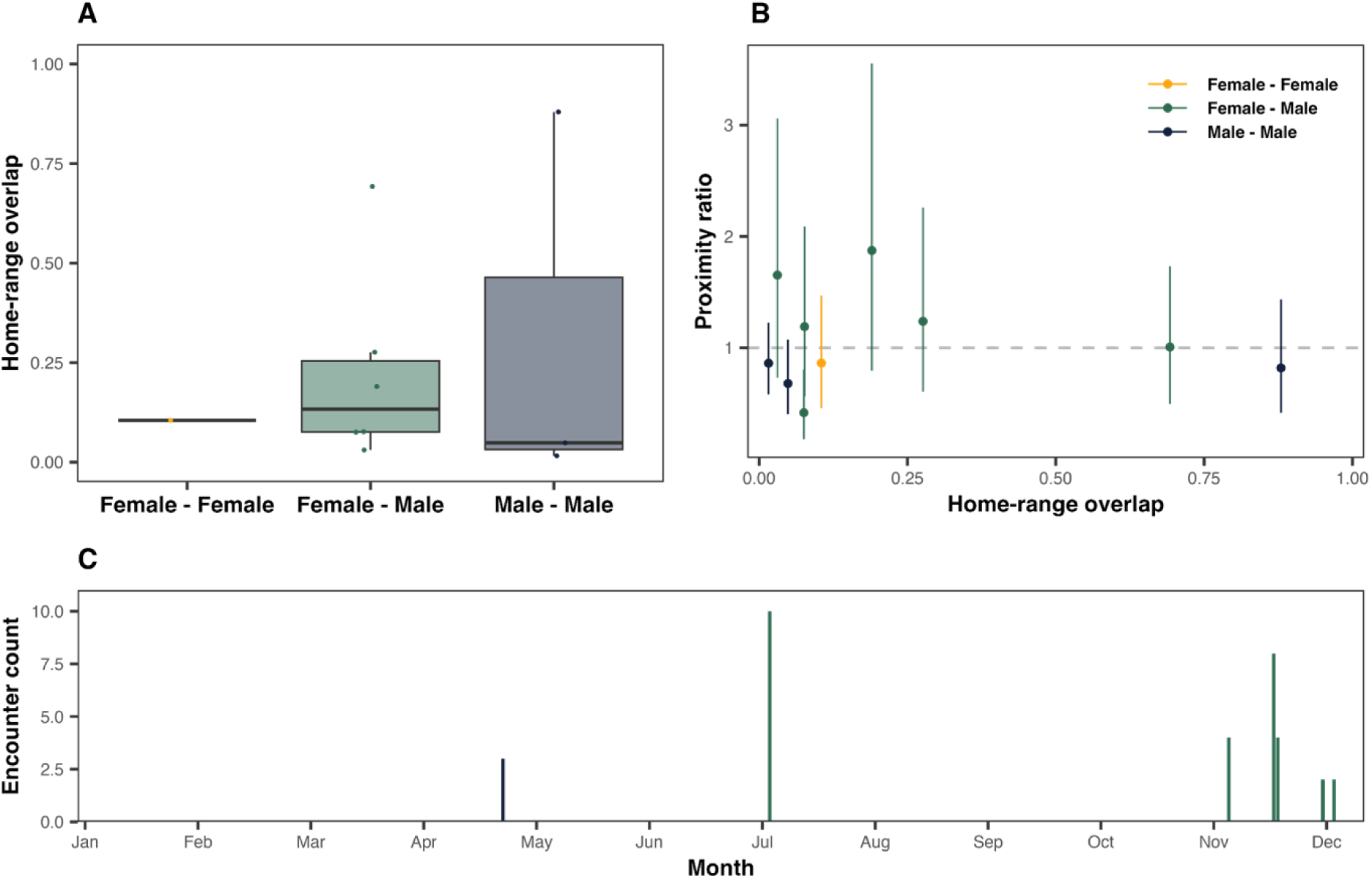
Patterns of jaguar socio-spatial behaviour. The boxplots in panel (A) depict the distribution of the home-range overlap estimates between the different pairwise combinations of sexes (n = 10 dyads). Panel (B) shows the estimated proximity ratios (+/- 95%CIs) as a function of home-range overlap for the 10 dyads. A proximity ratio of 1 indicates statistically independent movement, whereas values above 1 individuals were further apart than expected by independent movement, and vice versa for proximity ratios below 1. Panel (C) shows the distribution of encounter events over the annual cycle.

### Movement speed and activity patterns

On average, the monitored jaguars moved at a speed of 7.30 km/day (95% CI: 5.81 – 9.06 km/day), ranging from 3.92 km/day (95% CI: 3.85 – 3.99 km/day) to 10.36 km/day (95% CI: 10.26 – 10.45 km/day). We estimated the male/female ratio of movement speeds to be 1.09 (0.65–1.64), which includes 1, indicating no significant difference (Fig. 3A). There were also no significant relationships between movement speed and weight (β < -0.001; 95% CI: -0.003 – 0.002; Fig. 3B), nor age (β = -0.0015; 95% CI: -0.022 – 0.019; Fig. 3C). The jaguars on Maracá-Jipioca did, however, modulate their activity as a function of habitat. Animals were stationary 78.7% of the time, and moving the remaining 21.3%, but their probability of moving differed significantly between habitats (Λ_GLRT_ = 662.3, P < 0.001; Fig. 3D, Table 1). On average, the monitored animals tended to have a higher probability of moving in grasslands, water, and wetlands, versus a greater probability of resting while in forests and mangroves.

**Figure 3.**
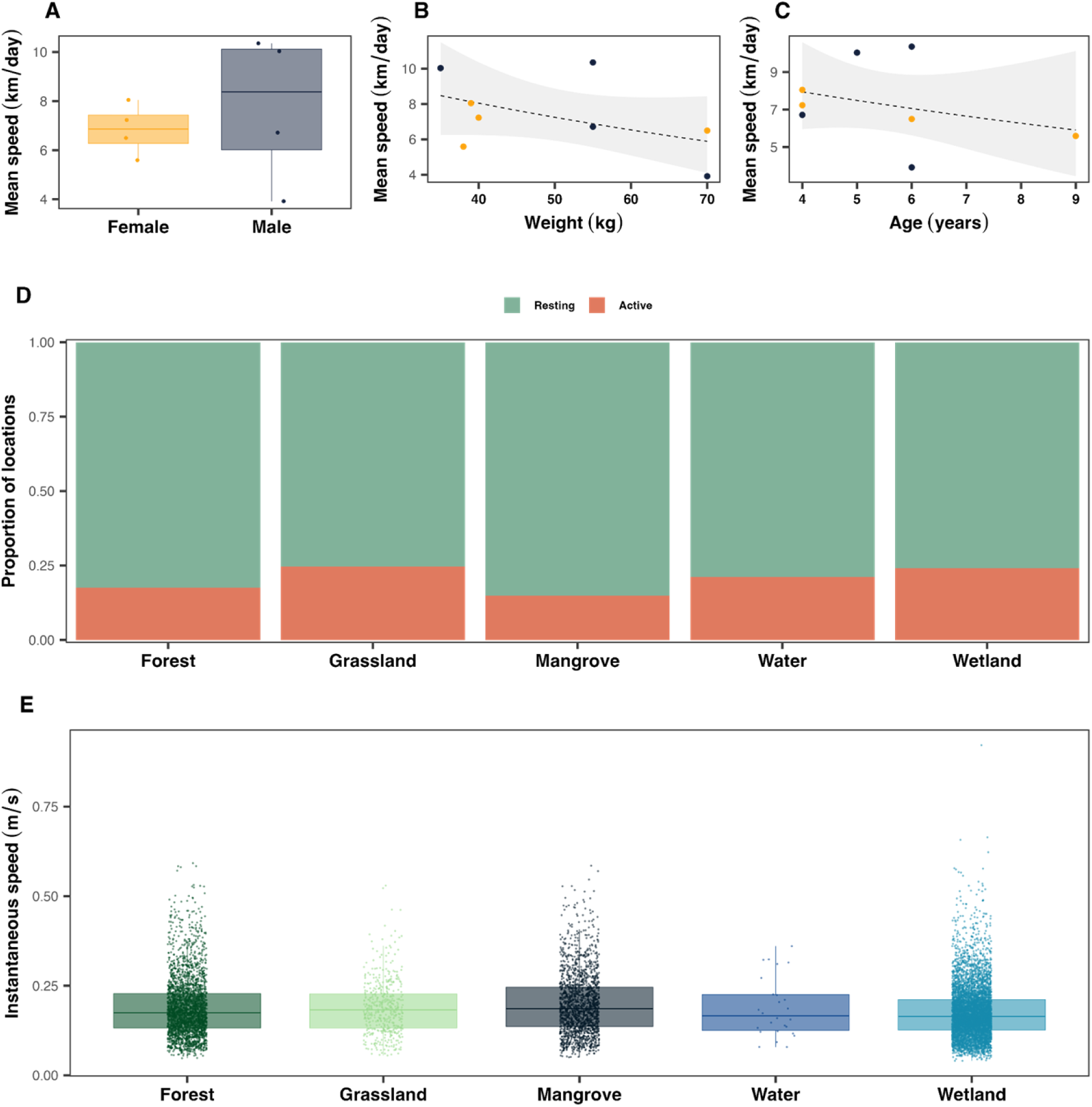
Jaguar movement speed and activity. Panel (A) depicts boxplots of movement speed between the sexes; as well as scatterplot of movement speed versus (B) weight, and (C) age. Panel (D) shows the distribution of locations classified of resting versus moving, based on the estimated movement speeds in each of Maraca’s five core habitat types. Panel (E) shows the eight monitored jaguars’ estimated instantaneous movement speeds while active in each of Maraca’s five core habitat types.

**Table 1.** The model estimated probabilities of a jaguar moving (P(moving)), and their movement speeds, along with 95% confidence intervals in each of Maraca’s five core habitat types.

|  | Forest | Grassland | Mangrove | Water | Wetland |
| --- | --- | --- | --- | --- | --- |
| **P(moving)** | 0.17<br>(0.14 - 0.21) | 0.24<br>(0.20 - 0.30) | 0.14<br>(0.11 - 0.17) | 0.24<br>(0.17 - 0.35) | 0.25<br>(0.21 - 0.30) |
| **Speed**<br>**(m/s)** | 0.177<br>(0.15 - 0.21) | 0.184<br>(0.16 - 0.22) | 0.171<br>(0.14 - 0.20) | 0.202<br>(0.16 - 0.25) | 0.182<br>(0.15 - 0.22) |

When active, the movement speed of the monitored jaguars also differed as a function of habitat (Λ_GLRT_ = 5.19, P < 0.001; Fig. 3E, Table 1). Jaguars’ average instantaneous movement speed was estimated as 0.18 m/s (range: 0.04 – 0.92 m/s). Compared to forests, these jaguars tended to move more quickly in grassland (β = 0.44, P < 0.001), wetlands (β = 0.49, P < 0.001), and water (β = 0.48, P = 0.034), but more slowly while moving through mangroves (β = -0.28, P < 0.0001).

Across all animals, activity peaked between 10:00 and 20:00 hrs (Fig. 4A). Interestingly, however, there were marked differences in the circadian rhythms between males and females, and a hierarchical GAM predicting activity that included an interactive effect between the time of day and sex was a significantly better match to the data than one that included time of day alone (Λ_GLRT_ = 836.3, P < 0.001). The patterns of this interaction term indicated that males and females avoided being active at the same time of day (Fig. 4B). The net result was that females tended to exhibit more diurnal activity, whereas males were active throughout the day, but with reduced daytime activity compared to females, and more pronounced crepuscular behaviour (Fig. 4C).

**Figure 4.**
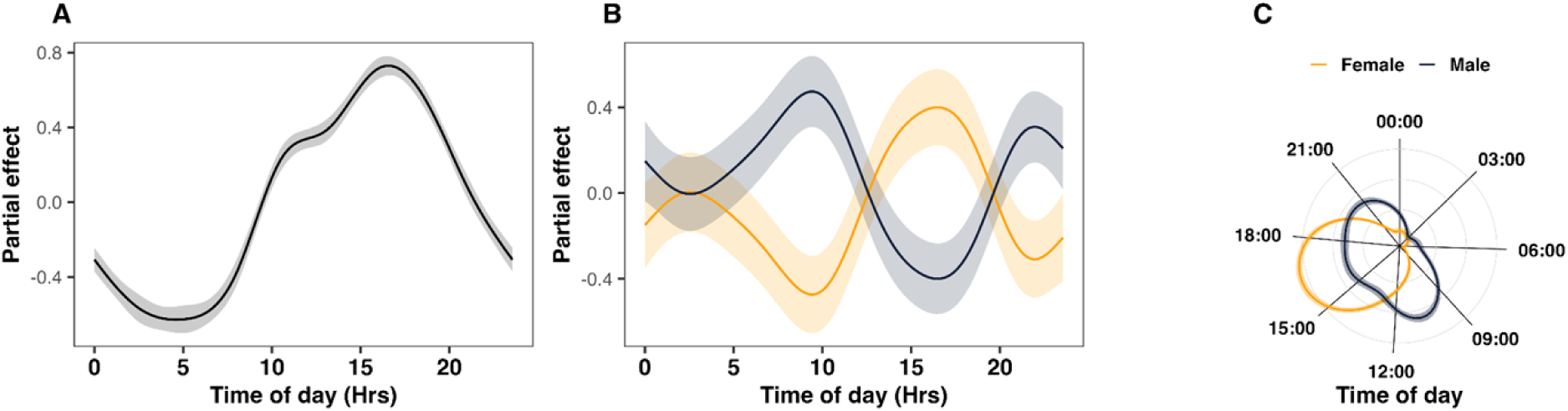
Circadian rhythms of jaguars on Maracá-Jipioca Islands. Panels (A) and (B) show the partial effects of a hierarchical generalised additive model fitted to the activity data. The curves in (C) show the full model predictions, demonstrating how the activity patterns of male and female jaguars change throughout a 24-hour period.

### Habitat use and selection

In addition to habitat dependent patterns in movement speed and activity, Maracá-Jipioca jaguars exhibited strong patterns of habitat preference (Fig. 5A). In general, individuals preferentially selected for grasslands, and forests, followed by mangroves, with comparatively low selection for the island’s wetlands. However, their habitat selection was influenced heavily by the interactive effect of distance to coast, with a strong preference for habitats ca. 100m from the coast (Fig. 5B).

**Figure 5.**
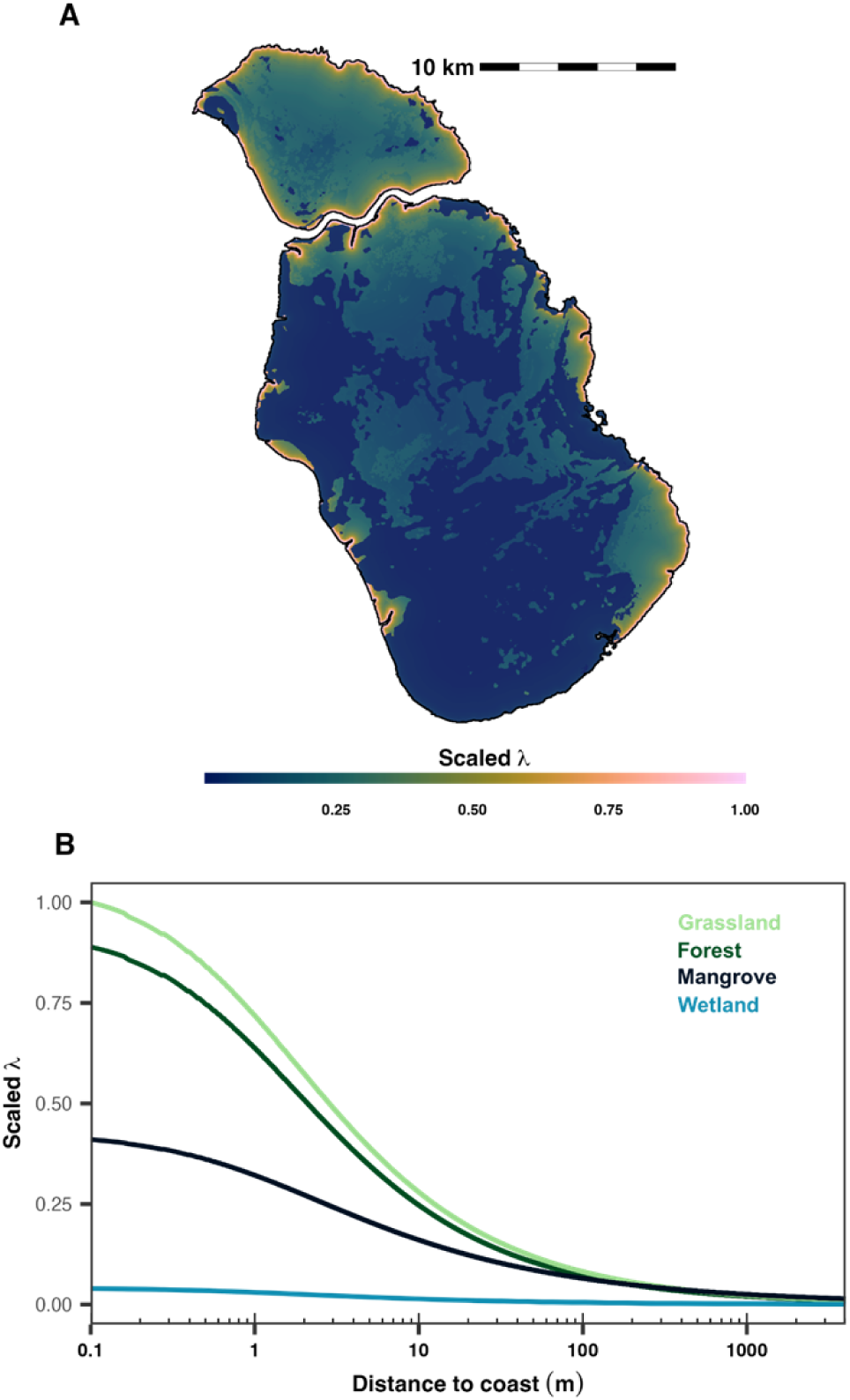
Jaguar habitat selection on Maracá-Jipioca Islands. Panel (A) shows the estimated habitat selection (λ) for jaguars on Maracá-Jipioca Islands. Panel (B) shows the estimated λ for each of the core habitat types as a function of distance to coast.

Jaguars on Maracá-Jipioca exhibited a strong preference for the Islands’ coastal habitats, and spent a substantial portion of their time directly at or near the island’s coast (Fig. 1A). Interestingly in this regard, we found that individuals adjusted their distance to the coast as a function of the tide, such that they tended to remain closer to the coast when the tide was low (β = 1.15, P < 0.001; Fig. 6). Although there was substantial residual variance (Fig. 6C), the model that included sea level was a significantly better predictor of jaguars’ distance to the coast than the null model that included only individual differences in behaviour (Λ_GLRT_ = 2999.2, P < 0.001).

**Figure 6.**
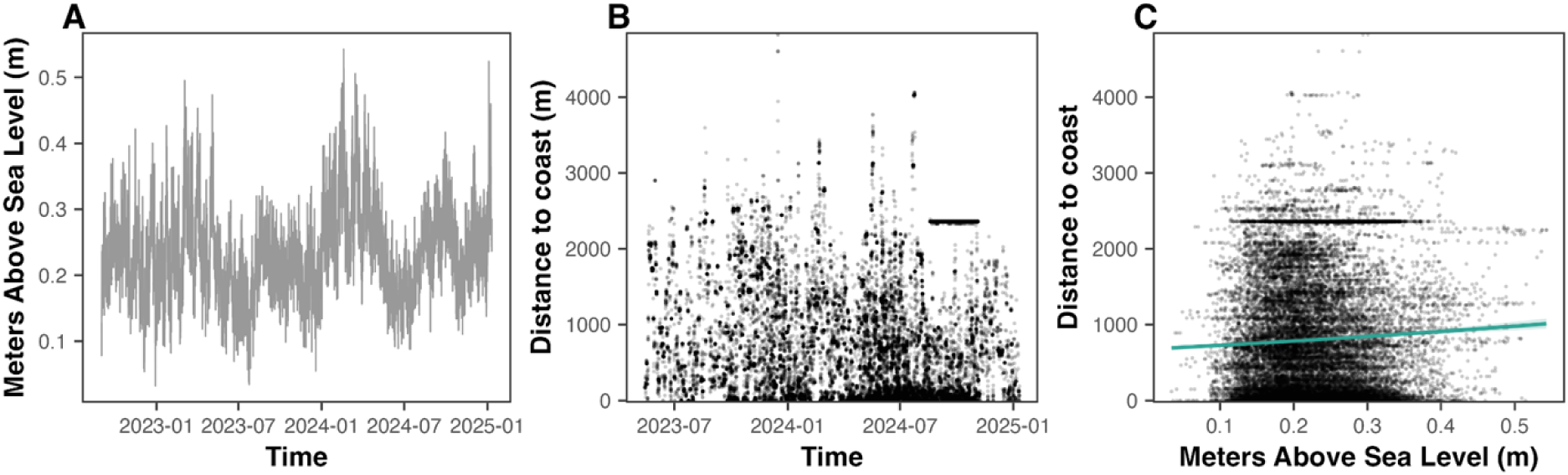
Jaguar habitat use and tidal patterns. Panel (A) shows the sea level, in meters above sea level, directly off of the coast of Maracá-Jipioca Islands between 2022-09-01 and 2025-01-10. The scatterplot in (B) shows the jaguars’ distance to the coast over the same time window. Panel (C) depicts the jaguars’ distance to coast as a function of the seal level. The green line shows the fitted generalised linear regression model.

## Discussion

This study offers the first detailed insight into the movement ecology of jaguars inhabiting an isolated Amazonian island system. Despite the spatial constraints imposed by the Maracá- Jipioca Ecological Station, jaguars maintained spatial and behavioural patterns similar with those observed in mainland populations, while also exhibiting unique adaptations to their insular environment. Unlike many previous studies conducted in human-altered landscapes, Maracá-Jipioca presents a rare example of a low-disturbance ecosystem (HFI ≈ 0), offering a valuable baseline for evaluating jaguar spatial ecology in the absence of anthropogenic pressures [13,56].

As with most continental jaguar populations [21,56], males on Maracá-Jipioca had significantly larger home ranges than females. While this pattern is consistent with jaguars’ known sex-specific space use, the absolute home-range sizes recorded on Maracá-Jipioca are remarkably small compared to those documented in other biomes. Here, the average home-range size across all monitored individuals was 51.9 km², with males averaging 77.8 km² and females just 19.5 km². In contrast, in the Amazon mainland biome, for example, males had an average home-range size of 211.6 km², versus 68.4 km² for females. In the Atlantic Forest, male and female ranges can exceed 460 km² and 260 km², respectively. Jaguars in the Pantanal, a prey-rich floodplain, still maintain larger average ranges than those on Maracá-Jipioca, with males spanning up to 152.4 km² and females up to 52.0 km². The Cerrado, with its open savannah landscapes, features some of the largest reported ranges, with individual males reaching 1,268.6 km² [21]. Recent meta-analyses confirm that home-range size increases with road density and decreases with forest cover and net primary productivity (NPP), with degraded areas demanding larger ranges for males in particular [56]. These contrasts suggest that spatial constraints imposed by island geography on Maracá-Jipioca are shaping reduced home range sizes, likely facilitated by adequate prey availability and the absence of human disturbance. Indeed, Alvarenga et al. (2025) and Morant et al. (2025) demonstrated that highly productive, roadless floodplain systems can sustain higher-than-average jaguar densities with correspondingly compact ranges [57,58].

Despite the limited area on Maracá-Jipioca, overlap between home ranges remained low (mean overlap: 0.24). Although this was comparable to the amount of overlap observed in a comparatively social population of jaguars in the Pantanal that specialized on aquatic prey [29], direct encounters between individuals were rare. Only 33 encounter events were detected during the entire tracking period, with the majority involving male–female pairs and only three between males. Notably, these male–male encounters occurred between a single pair of individuals with the highest home-range overlap, suggesting that while spatial tolerance may increase under aquatic prey bases [29] restricted or high-density conditions, direct interactions may still remain infrequent. This pattern contrasts with findings from the Pantanal, where radiocollared males have shown extensive home-range overlap and were located within 200 m of each other more frequently than expected by chance, indicating some degree of spatial tolerance or sociality [59]. However, even in that context, social interactions remained limited and did not deviate substantially from the species’ broadly solitary social structure. That jaguars on Maracá-Jipioca maintain low overlap and rare direct contact, even in a flooded, resource-accessible, and spatially restricted setting, shows their capacity to preserve territorial spacing. Such spacing behaviour is consistent with optimal territory partitioning observed in high-quality habitats with minimal human risk, where competition is mitigated by abundant resources [58]. However, it is also possible that not all animals in the area were captured and equipped with GPS devices, so our findings in this regard should be interpreted with caution.

We estimated the average daily movement on Maracá-Jipioca Island to be 7.3 km/day, with no significant differences observed between males and females or across age and body size classes. These values are consistent with the upper range of those observed in other Brazilian biomes. In the Amazon, for example, average daily travel for GPS-tracked jaguars ranged from 2.3 to 5.9 km/day, in the Atlantic Forest, movement rates were generally higher, reaching up to 15.4 km/day for males and 11.5 km/day for females [21]. In the Pantanal, daily travel distances varied widely, with some individuals traveling 8.9–11.4 km/day, while others showed slower movement at 4.0 km/day [21]. These data place Maracá-Jipioca jaguars well within the ecological norm, especially relative to fast-moving individuals from the Atlantic Forest and Pantanal. Morant et al. (2025) found that monthly displacement tends to increase with human footprint (up to HFI ≈ 5.6), consistent with our observation that low-disturbance conditions on Maracá-Jipioca yield moderate, stable movement rates. Despite similarities in mean speed, movement behaviour on Maracá-Jipioca was highly habitat-specific. Jaguars moved significantly faster in open habitats such as grasslands, wetlands, and even flooded areas, while slowing substantially in forests and mangroves. This contrasts with Amazonian jaguars, who primarily traverse dense forest and exhibit more constrained movement (mean: 2.3–5.9 km/day), and more closely resembles the broader and more dynamic movements of Pantanal jaguars inhabiting savannah-like mosaics [21]. While the ecosystem composition, and insular nature of Maracá-Jipioca may be unique, the consistent finding across all regions appears to be that habitat structure, and not sex or age, is the key determinant of jaguar movement patterns. Open habitats likely promote faster, more exploratory movement in search of prey, while dense vegetation facilitates ambush hunting and cover-based travel. Maracá-Jipioca’s spatial heterogeneity and productivity, combined with its flooded margins, likely create a mosaic that supports efficient prey tracking without requiring large-fast-scale displacements.

Interestingly, the activity patterns of jaguars on Maracá-Jipioca were notably distinct from those described in most continental populations. Across all individuals, activity peaked between 10:00 and 20:00, with a striking degree of diurnality. Females were particularly active during the day, whereas males exhibited a more crepuscular rhythm, suggesting a potential temporal partitioning between the sexes. In contrast, jaguars in mainland ecosystems, such as the Pantanal and Atlantic Forest, are predominantly crepuscular or nocturnal, with significantly reduced daytime activity [21,59]. These patterns are thought to reflect behavioural adaptation to avoid human presence [60], which is largely absent from Maracá-Jipioca. Indeed, studies from Belize and other human-impacted regions have shown strong nocturnal tendencies in jaguars associated with anthropogenic disturbance [61,62]. Morant et al. (2025) found that male diurnality declines sharply when HFI exceeds 3.3, reinforcing the idea that jaguars on Maracá-Jipioca maintain more natural activity rhythms under negligible disturbance. However, the marked diurnality may also be linked to hunting behaviour [6] and the islands’ unique prey base and ecological context. Jaguars on Maracá-Jipioca are known to consume a wide variety of prey, including aquatic and semi-aquatic species such as fish, caimans, capybaras, and even marine dolphins, as documented by Dias et al. (2023) [22,63]. This diet, heavily reliant on visually detected and aquatic prey, could favour daylight hunting when visibility is higher. Moreover, the observed sexual differentiation in activity, where males exhibit more crepuscular behaviour while females are more active during the day, may reflect differing reproductive roles, prey partitioning, or energetic needs. Such patterns have rarely been documented in mainland populations, underscoring the potential influence of island-specific ecological pressures. Further behavioural and dietary studies are needed to disentangle the drivers of these novel temporal activity patterns and to assess how prey type, detection mode, and habitat use interact to shape jaguar rhythms in insular environments.

Interestingly, jaguars on Maracá-Jipioca spent a substantial proportion of their time directly at or near the island’s coast and exhibited a measurable response to tidal variation. Specifically, individuals tended to stay closer to the coast during low tide, suggesting that their spatial use is at least partially influenced by hydrological cycles. This behaviour may reflect strategic foraging, as many of their key prey species, such as fish, caimans, and capybaras, are more accessible or concentrated in shallow waters during low tide. While substantial individual variability was observed, the inclusion of sea level significantly improved model predictions of coastal proximity, indicating a functional link between tidal state and jaguar movement. These findings suggest that, in this insular Amazonian system, jaguars modulate their coastal behaviour in response to environmental rhythms, possibly to enhance prey encounter rates. The observed fidelity to coastal areas, often within 100 m of the waterline, mirrors the spatial optimisation seen in other flooded, high-NPP systems where aquatic prey resources are predictably structured [57]. Future studies integrating finer-scale movement data and prey availability across tidal phases would help clarify the ecological mechanisms underlying this pattern.

In summary, Maracá-Jipioca represents a natural laboratory for understanding jaguar ecology under near-pristine conditions. The absence of direct human pressure, coupled with high productivity and aquatic prey abundance, supports reduced home ranges, slower movement, and higher diurnality. These traits contrast with the expanded ranges and altered behaviours reported from disturbed systems, highlighting the role of habitat integrity in shaping large carnivore ecology [56,58]. Yet, the total land area available (ca. 600 km^2^) remains well below the ∼1,700 km² required for a genetically viable population under typical conditions [21], raising questions about long-term persistence. Ongoing demographic, genetic, and landscape-scale monitoring will be essential to determine whether Maracá-Jipioca acts as a demographic source, sink, or closed system, and how it may respond to climate-driven changes in sea level and habitat structure.

## Acknowledgments

We would like to acknowledge the many field and laboratory technicians who assisted with the data collection. We thank ICMBIO for giving the permits to carry this study.

## Author contributions

Conceptualization: LS, FBLP, MAMMF, MJN, TJS, ATAJ, GRA

Methodology: MJN, LS, RAM

Investigation: MJN

Visualization: MAMMF, MJN

Funding acquisition: MJN, ATAJ

Project administration: LS, MJN, ATAJ

Supervision: LS, MJN

Writing – original draft: LS, FBLP, MAMMF, MJN

Writing – review & editing: all authors

## Data and materials availability

The code and data required to reproduce all of the analyses and figures in the manuscript are openly available at: https://github.com/QuantitativeEcologyLab/island_jaguars. The MapBiomas habitat classification rasters are all openly available in publicly accessible repositories.

## Supplementary Materials

**Table S1.** Detailed description of the seven jaguars monitored with GPS collars in Maracá-Jipioca. One of the females,

| Campaign | Name | Colar | Sex | Age | Weight | Start | End | Interval | Duration | Points |
| --- | --- | --- | --- | --- | --- | --- | --- | --- | --- | --- |
| 1 | Iemanja1 | 707821 | Female | 4 | 39 | 2019-10-04 | 2020-08-09 | 30 | 10.47 | 12,897 |
| 1 | Iara | 704795 | Female | 4 | 40 | 2019-10-09 | 2020-10-22 | 30 | 12.80 | 16,277 |
| 1 | Netuno | 704741 | Male | 6 | 55 | 2019-10-19 | 2020-10-08 | 30 | 12.03 | 14,988 |
| 2 | Iemanja2* | 696435 | Female | 9 | 38 | 2023-05-13 | 2024-07-06 | 60 | 14.19 | 7,010 |
| 2 | Iranildo | 696417 | Male | 4 | 55 | 2023-09-24 | 2024-11-04 | 60 | 13.77 | 8,260 |
| 3 | ID696490B | 696490 | Male | 6 | 70 | 2024-04-19 | 2024-11-04 | 60 | 6.73 | 3,775 |
| 3 | ID717047B | 717047 | Female | 5 | 35 | 2024-05-03 | 2024-07-16 | 30 | 2.49 | 1,478 |
| 3 | ID696469B | 696469 | Male | 6 | 70 | 2024-05-15 | 2025-01-24 | 60 | 8.59 | 4,249 |

**Figure S1.**
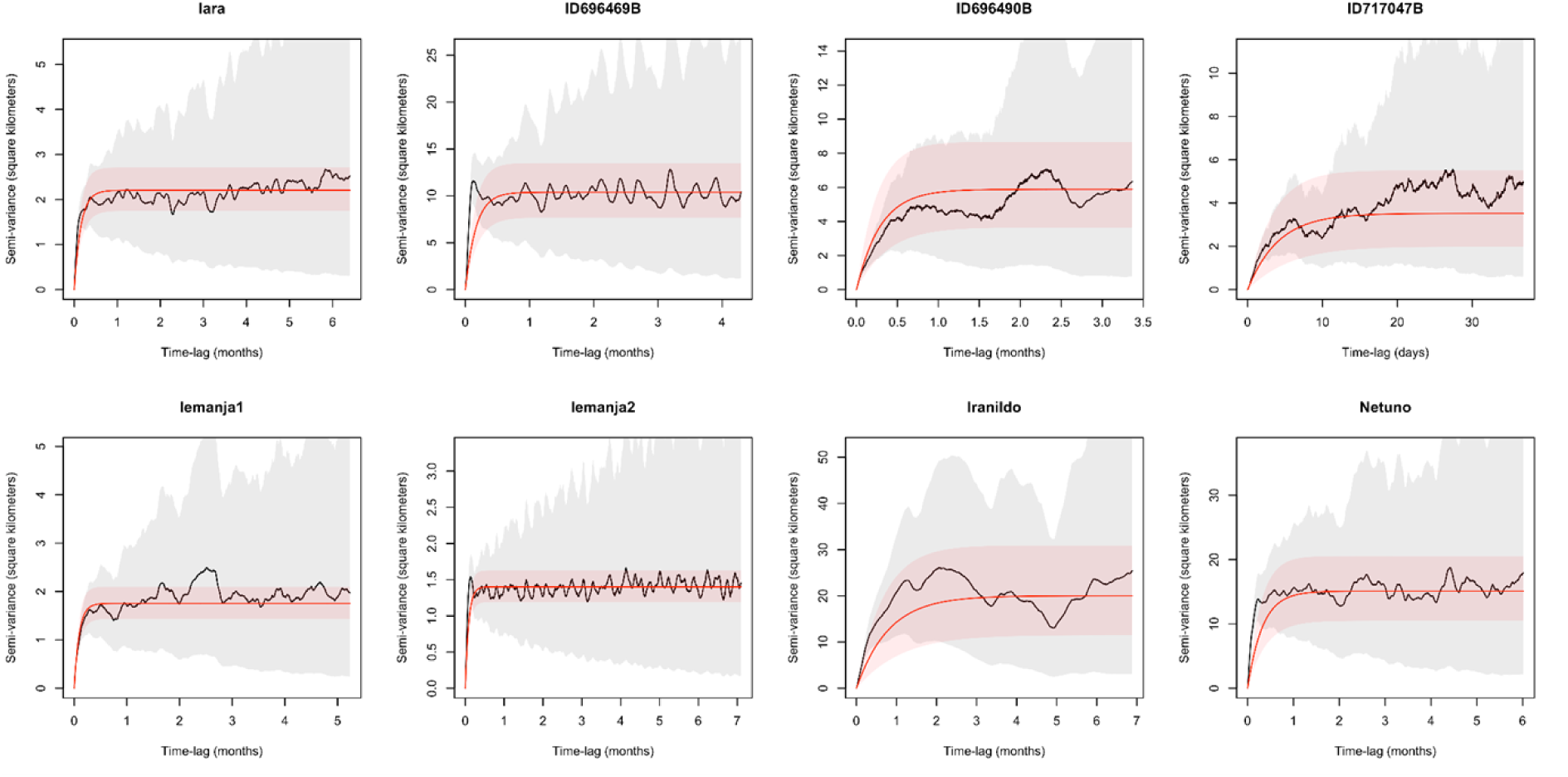
Semi-variograms and fitted movement models. The empirical semi-variograms (black lines) and 95% confidence intervals (grey shading), as well as the fitted movement models (red lines) and 95% confidence intervals (red shading) for each jaguar. Not how all of the empirical variograms have clear asymptotes, indicating range-residency, and the fitted models accurately capture trends in the data.

